# Structure of a germline-like human antibody defines a neutralizing epitope on the SARS-CoV-2 spike NTD

**DOI:** 10.1101/2021.07.08.451649

**Authors:** Clara G. Altomare, Daniel C. Adelsberg, Juan Manuel Carreno, Iden A. Sapse, Fatima Amanat, Ali H. Ellebedy, Viviana Simon, Florian Krammer, Goran Bajic

## Abstract

Structural characterization of infection- and vaccination-elicited antibodies in complex with antigen provides insight into the evolutionary arms race between the host and the pathogen and informs rational vaccine immunogen design. We isolated a germline-like monoclonal antibody (mAb) from plasmablasts activated upon mRNA vaccination against SARS-CoV-2 and determined its structure in complex with the spike glycoprotein by cryo-EM. We show that the mAb engages a previously uncharacterized neutralizing epitope on the spike N-terminal domain (NTD). The high-resolution structure reveals details of the intermolecular interactions and shows that the mAb inserts its HCDR3 loop into a hydrophobic NTD cavity previously shown to bind a heme metabolite, biliverdin. We demonstrate direct competition with biliverdin and that - because of the conserved nature of the epitope – the mAb maintains binding to viral variants B.1.1.7 and B.1.351. Our study illustrates the feasibility of targeting the NTD to achieve broad neutralization against SARS-CoV-2 variants.

## Introduction

Severe acute respiratory syndrome coronavirus 2 (SARS-CoV-2) has officially caused more than 185 infections and more than 4 million official deaths worldwide (World Health Organization). Immune responses mounted upon COVID-19 (Coronavirus disease 2019) have been a subject of active investigations by many groups. As safe and effective vaccines are developed and administered in record time^1,2^ there is an urgent need to better understand the quality of the vaccine-induced immune responses, the broadly neutralizing epitopes targeted and their effectiveness against newly emerging, potentially more transmissible viral variants. Understanding the immunodominance landscape of the major antibody target – the spike glycoprotein – at a structural level will identify the requirements for broader SARS-CoV-2 antibody responses and provide the foundation for developing the next generation of vaccines.

The viral spike glycoprotein is both the attachment factor which binds angiotensin converting enzyme 2 (ACE2) on host cells and the viral fusogen that mediates viral membrane fusion with that of the host cell^3^. The fusion step depends on a furin-mediated cleavage, resulting in the generation of N-terminal S1 and C-terminal S2 domain^4^. The second, subsequent cleavage of S2 is mediated by a serine protease, TMPRSS2 or by cathepsins^5^. The spike glycoprotein is the main target of neutralizing antibody responses and hence the focus of most vaccines. Antibody responses to natural infection in the serum, memory B cell compartment, and to a lesser degree at mucosal surfaces, against spike have been well characterized in terms of kinetics, binding specificity and neutralization potency^6-18^. Anti-SARS-CoV-2 spike serum antibody titers after natural infection are variable, may decline to some degree over time^19,20^ and have suboptimal neutralization activity against more recent viral variants, despite being protective^21,22^. Antibodies derived from memory B cells target both unique, and to a certain extent, also overlapping epitopes that contribute to polyclonal epitopic coverage of spike and ensure preserved binding to viral variants of concern^23-30^. We have recently shown that immunization with mRNA vaccines results in antibodies targeting not only the receptor binding domain but also the N-terminal domain (NTD)^31^.

We focus here on the early events of B cell activation after SARS-CoV-2 vaccination to structurally profile novel antibody epitopes. We previously identified a neutralizing monoclonal antibody (mAb), PVI.V6-14, derived from the plasmablast response mounted by a naïve study participant after two doses of mRNA vaccine whose heavy and light chains both contained no somatic hypermutation^31^. We determined its high-resolution structure in complex with the SARS-CoV-2 spike by single-particle electron cryomicroscopy (cryo-EM) and showed that it bound a lateral side of the N-terminal domain (NTD). The interaction was mainly mediated by the heavy complementarity-determining region 3 (HCDR3) loop with minimal contacts from the light chain. We found that mAb PVI.V6-14 belongs to an as of yet undescribed class of antibodies that bind within a hydrophobic cavity, previously identified to bind a heme metabolite, biliverdin^32^. Our functional binding and neutralization data confirm that the antibody competes with biliverdin and also underscores the antibody’s capacity to recognize emerging viral variants of concern. Our study puts forward a concept for a therapeutic combination antibody cocktail that comprises both receptor binding domain (RBD) as well as NTD neutralizing antibodies. Collectively our results inform on the rational design of a novel class of immunogens for next generation vaccines that provide broad protection against currently circulating as well as future, newly emerging SARS-CoV-2 variants of concern.

## Results

### Neutralizing antibody PVI.V6-14 binds a lateral cavity in the NTD

We previously reported on the isolation and characterization of plasmablast-derived antibodies from individuals who received the Pfizer/BioNTech mRNA SARS-CoV-2 vaccine BNT162b2^31^. We noted that the overall neutralizing antibody responses were directed towards both the RBD and NTD indicating co-dominance of these two spike domains. Since NTD emerges to be an important component of the vaccine-induced responses, we wanted to expand on our understanding of neutralizing epitopes in this region of the SARS-CoV-2 spike for which only limited structural information is currently available. We therefore focused our attention on participant V6 who mounted a strong neutralizing antibody response to the NTD region of spike upon vaccination. We selected, for further analysis, a neutralizing antibody, PVI.V6-14 that bound to NTD and whose V(D)J sequence was identical to the germline. Indeed, PVI.V6-14’s heavy chain variable (V_H_) sequence is identical to the human IGHV4-39*01 germline sequence (Figure S1a). The rearranged V(D)J complementarity determining region (CDR) 3, that encodes 21 amino-acid residues, is composed of IGHD3-10*01 and IGHJ4*02 (Figure S1b). The kappa light chain too was unmutated and encoded by IGHKV1-12*01 (Figure S1c-d). We determined its structure in complex with the spike glycoprotein to a high resolution using single-particle cryo-EM (Figure 1 and Figure S2). PVI.V6-14 binds NTD on the side that is anti-clockwise looking down on RBDs and perpendicular to the three-fold axis of the spike (Figure 1a-b). There are two Fabs bound per spike trimer in the final cryo-EM reconstruction. Incidentally, RBDs are in the “down” configuration on the two protomers whose NTDs are in complex with Fab; the unbound protomer has its RBD partially “up”. Our NTD-Fab focused, locally refined map shows that the heavy chain CDR3 loop protrudes deep into the NTD cavity (Figure 1c-d). PVI.V6-14 appears to stabilize NTD as the cryo-EM map shows clear, undisrupted volume for the entire domain allowing us to trace the complete polypeptide chain with high confidence (Figure 1d).

**Figure 1.**
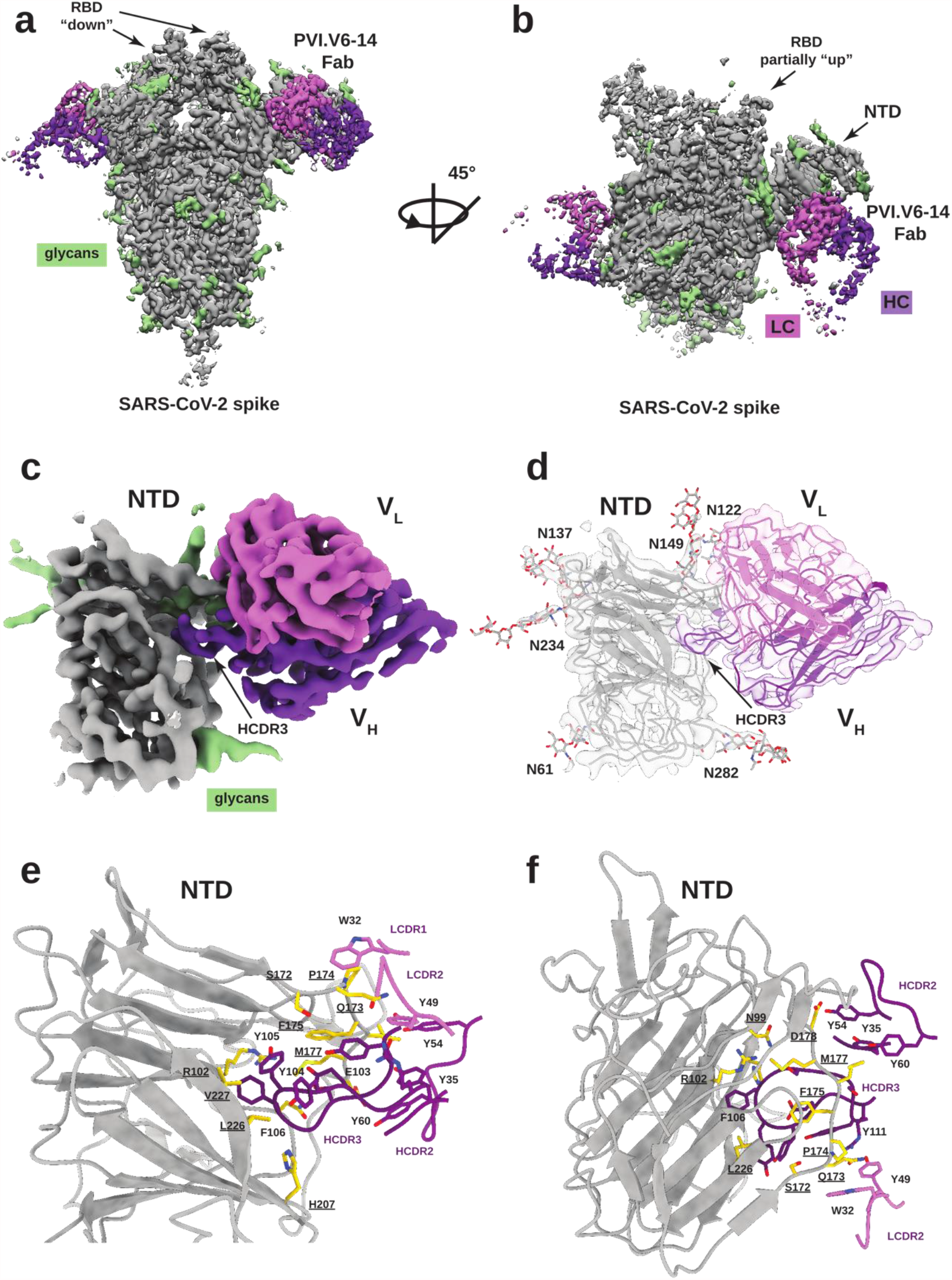
PVI.V6-14 mAb recognizes a novel epitope on the SARS-CoV-2 spike NTD. **(a-b)** Cryo-EM reconstruction of the SARS-CoV-2 spike trimer (grey, glycans in green) with PVI.V6-14 Fab (heavy chain in purple, light chain in pink) bound to the NTD, with two 45-degree rotated views. Two Fabs are bound, two RBDs are in the “down” conformation. **(c)** Focus refined map of PVI.V6-14 bound to the NTD at 3.2 Å nominal resolution, with HCDR3 inserted in the hydrophobic NTD pocket. **(d)** Atomic model built into the cryo-EM map of the PVI.V6-14:NTD complex. **(e-f)** Details of the intermolecular interactions between PVI.V6-14 and the NTD are dominated by the HCDR3 loop. NTD interacting amino-acid residues are shown in gold with residues number labels underscored.

PVI.V6-14 binds on the NTD between two glycans – N122 and N282 – and contacts the hydrophobic cavity primarily through its HCDR3 loop composed of a string of aromatic amino-acid residues Tyr104, Tyr105 and Phe106. In particular, Tyr105 interacts with Arg190, Phe192 and His207 in NTD. Additional HCDR3 contacts are provided by Glu103 and Ser108 that form hydrogen bond donor and acceptor, respectively, and interact with Gln173 and Pro174 (Figure 1e-f). The V_H_ makes additional contacts with NTD through HCDR1 Tyr35 and HCDR2 Tyr54 and Tyr60. Minor light chain contacts are supplied by the LCDR1 Trp32 that pi-pi stacks against Pro174 on NTD and LCDR2 Tyr49 which hydrogen bonds with Gln173 (Figure 1e-f).

### PVI.V6-14 binds viral variants of concern

We next performed a sequence alignment of the spike proteins of 35 different sarbecoviruses and noticed that the NTD cavity was comprised of amino-acid residues that were conserved across SARS-CoV-2 isolates and bat coronaviruses with pandemic potential (Figure S3). We hypothesized that PVI.V6-14 would also bind to the emerging variants of concern (VOC). Indeed, this antibody bound well to the spike glycoproteins of B.1.1.7 and B.1.351 (Figure 2a) but lost binding to the P.1 variant (Figure 2a). We next mapped the mutations of the viral variants of concern onto NTD region within the structure of our complex (Figure 2b). The structure supports the lack of impact of the substitutions and deletions specific for the B.1.1.7 (Alpha) and B.1.351 (Beta) lineages. It also highlights the relevance of the arginine residue at position 190, which is mutated to a serine in the P.1 (Gamma) lineages. In our previous study we noticed that PVI.V6-14 did not neutralize B.1.1.7 and B.1.351 viruses^31^. Our NTD structure, therefore, offers a molecular rationale for the antibody’s breadth and sensitivity to particular substitutions within NTD.

**Figure 2.**
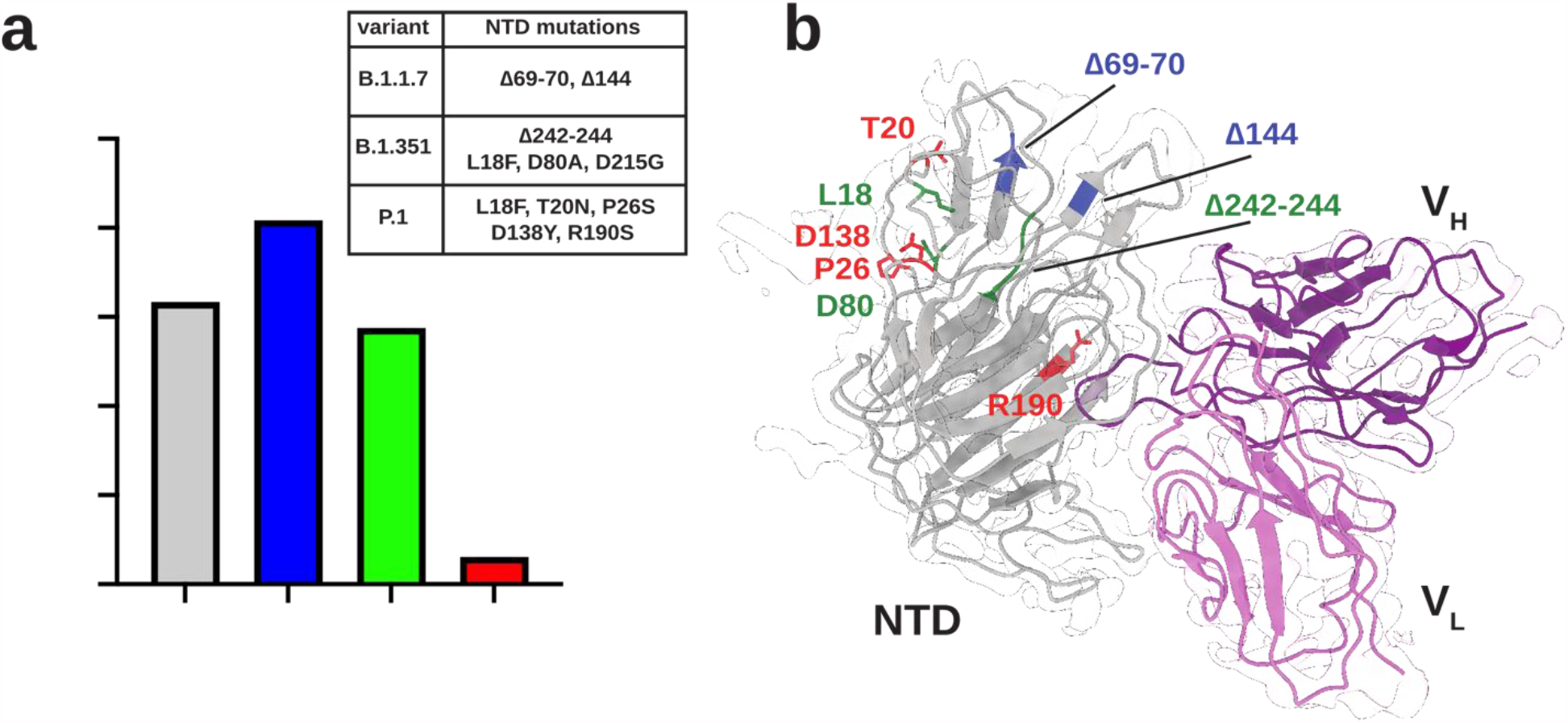
PVI.V6-14 binds viral variants of concern B.1.1.7 and B.1.351 but not P.1. **(a)** Binding of PVI.V6-14 IgG to HexaPro (WT;grey), B.1.1.7 (blue), B.1.351 (green) and P.1 (red) spikes of SARS-CoV-2. Area under the curve (AUC) was calculated by subtracting the average of blank values plus 3 times standard deviation of the blank values. Shown are means of a representative experiment performed in triplicates. Anti-polyhistidine IgG was used as positive control across the ELISA plates. The inset table recapitulates the VOC amino-acid mutations. **(b)** Structural mapping of the VOC mutations onto NTD in complex with PVI.V6-14 Fab. The structure explains the mAb dependence on R190 residue in NTD and the diminished binding to P.1 VOC. Color scheme same as in **(a)**.

### Antibody PVI.V6-14 directly competes with and is inhibited by biliverdin

A previous study reported the structure of a heme metabolite, biliverdin, bound within the hydrophobic NTD pocket^32^. We superposed the structure of our Fab bound NTD with that of the biliverdin bound one and found that our antibody, through its HCDR3 aromatic amino-acid residues, was a molecular mimic of the tetrapyrrole molecule (Figure 3a). Indeed, the two complexes share most of the NTD contact residues. To corroborate this observation, we performed a bio-layer interferometry (BLI)-based competition assay with PIV.V6-14 and biliverdin (Figure 3b). We observed a concentration-dependent inhibition of PVI.V6-14 binding to recombinant NTD by biliverdin. We next asked if biliverdin could interfere with binding of this antibody class to a virus and performed neutralization assays on an authentic SARS-CoV-2 isolate (Figure 3c-d). We found that biliverdin abrogated PVI.V6-14 neutralization at high concentration (Figure 3c). The binding of an RBD antibody 2C08^33^ was unaffected (Figure 3c). A remdesivir control is shown in Figure 3d.

**Figure 3.**
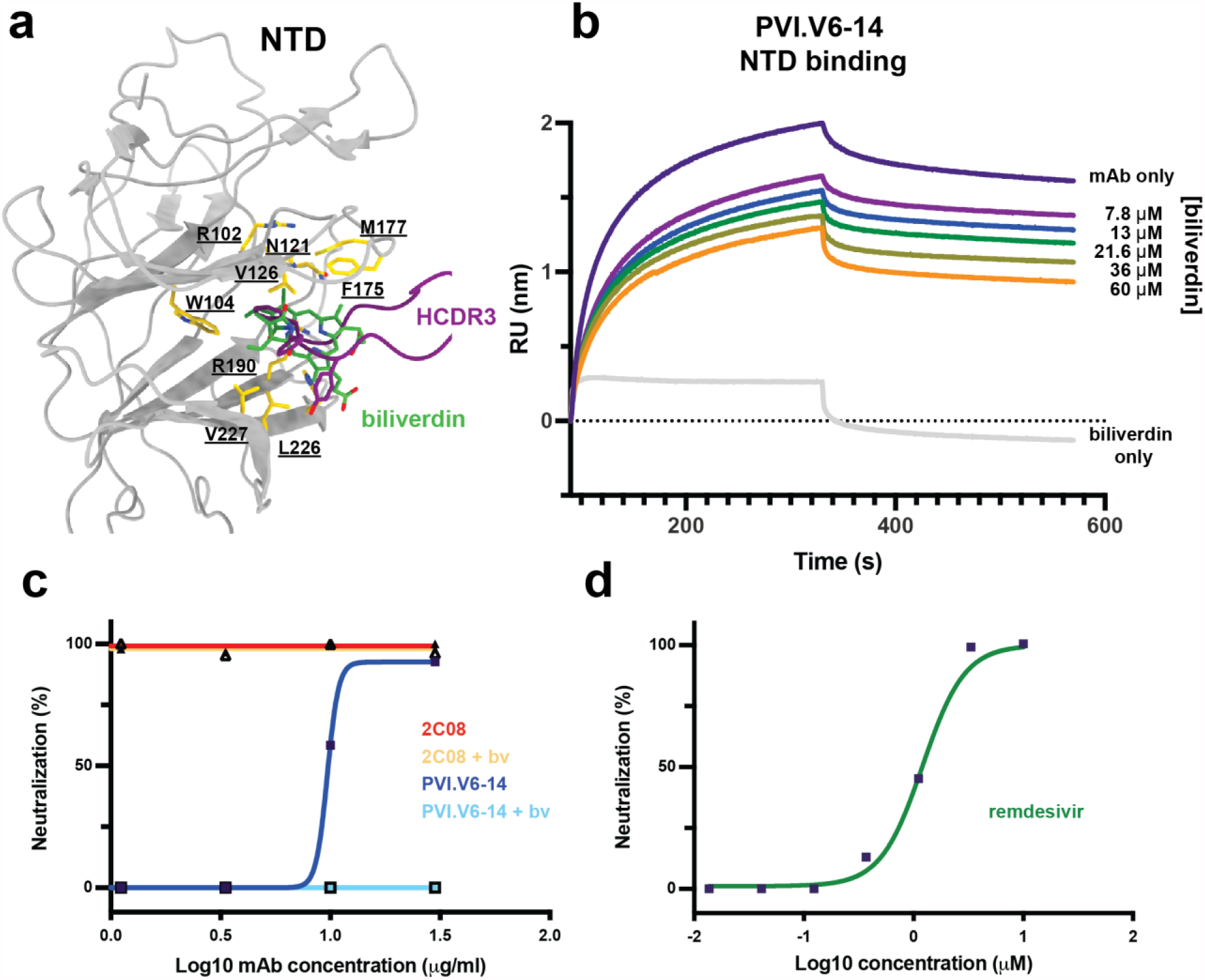
Biliverdin directly competes with PVI.V6-14. **(a)** Structural superposition of the biliverdin bound NTD (PDB ID 7B62) with the PVI.V6-14 bound NTD (this study, PDB ID 7RBU). Biliverdin is shown in green and PVI.V6-14 HCDR3 loop in purple sticks. NTD interacting amino-acid residues are shown in gold with residues number labels underscored. **(b)** Bio-layer interferometry (BLI)-based competition assay of biliverdin with PVI.V6-14 on recombinant NTD. **(c)** Neutralization assay with authentic SARS-CoV-2 virus, of NTD binding PVI.V6-14 and RBD-binding 2C08 mAb with and without biliverdin (bv). PVI.V6-14 directly competes with biliverdin while 2C08 neutralization activity is unaffected by biliverdin. **(d)** remdesivir neutralization control.

### PVI.V6-14 engages a distinct epitope

A recent study reported a structure of an NTD-targeting antibody, P008_056, that also competes with biliverdin for binding to NTD^32^. This antibody, however, does so allosterically, while PVI.V6-14 is a direct competitor. Indeed, the two antibodies approach the spike NTD from a different angle (Figure 4a). The binding of the two antibodies to NTD is, however, mutually exclusive for two reasons 1) there would be severe steric clash of the two V_H_ domains (Figure S4b) and 2) the NTD configuration must be “open” for P008_056 to bind and “closed” for PVI.V6-14 (Figure 4c). Indeed, the PVI.V6-14 bound NTD structure resembles more the biliverdin bound one, whereas the P008_056 complex induces a conformational change that reconfigures the 175-185 loop in a downward and out (Figure 4c).

**Figure 4.**
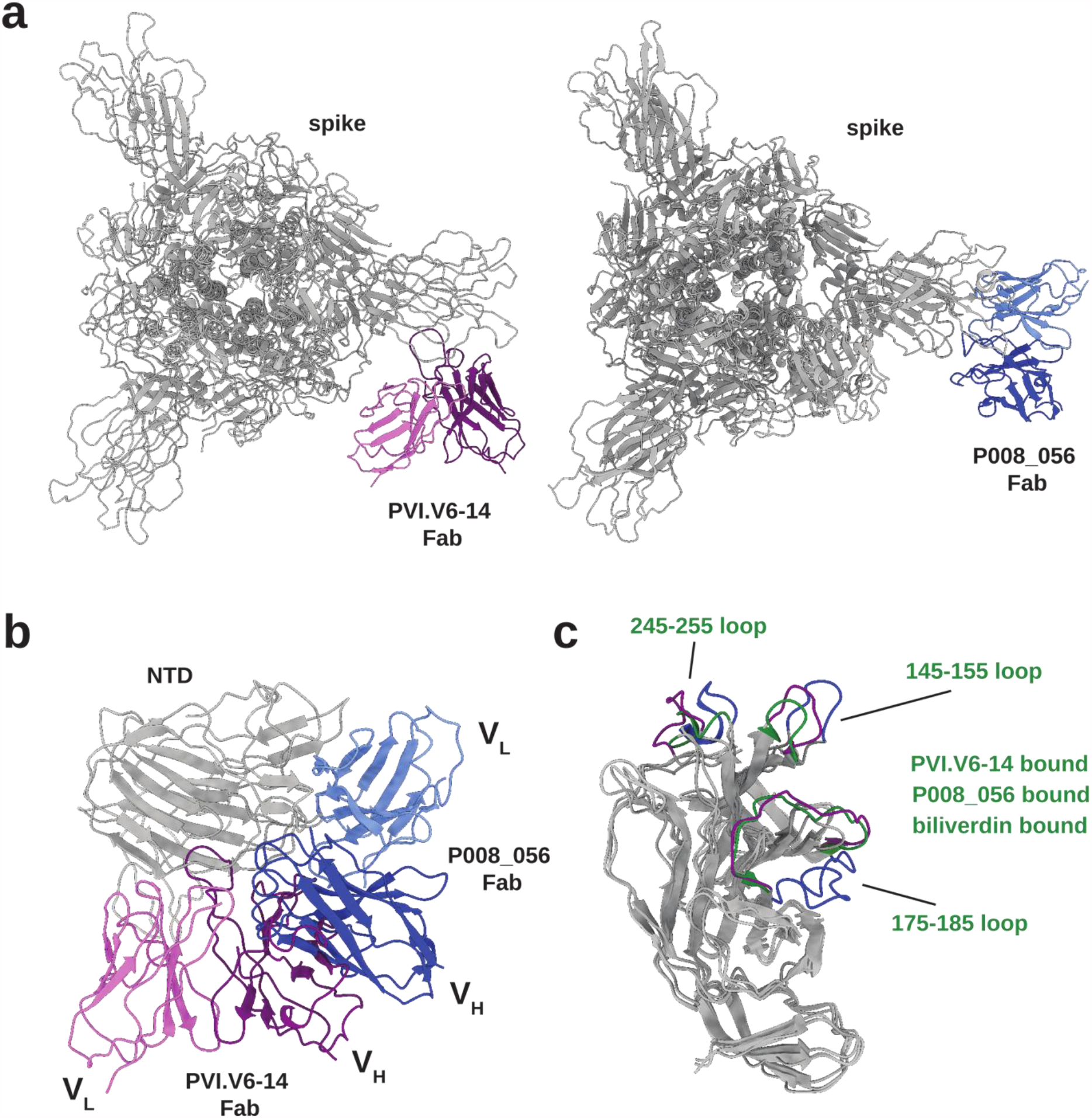
Structural comparison of PVI.V6-14 and P008_056. **(a)** Atomic models of the PVI.V6-14- and P008_056-bound SARS-CoV-2 spike. While both Fabs compete with biliverdin, their epitopes and angles of approach are different. (b) Structural superposition of PVI.V6-14 and P008_056 Fab:NTD complexes shows that the binding of the two classes of antibodies is mutually exclusive. (c) Structural superposition of the PVI.V6-14- (purple), P008_056- (blue) and biliverdin-bound (green) NTD. Major structural rearrangements are localized to the 245-255, 145-155 and 175-185 loops. PVI.V6-14 NTD conformation is similar to the biliverdin-bound one.

## Discussion

Establishment of specific and durable humoral immunity relies upon the activation of specific B-cell populations that respond to infection or vaccination. Plasmablasts are a part of early responses, and their immunoglobulin repertoire most likely reflects the composition of the early antigen-specific polyclonal serum. Understanding the specificities and the neutralization potential of those responses is therefore of paramount interest for infectious disease research and vaccinology. Additionally, the quality and the distribution of epitopes targeted may differ between natural infection and vaccination. We previously reported that infection with SARS-CoV-2 results in varying levels of antibody titers against the spike glycoprotein but that mRNA vaccination, in contrast, induces consistently high titers^31,34,35^. We also found that vaccine-derived antibody responses target more non-neutralizing epitopes than the responses from COVID-19 survivors^31^. We therefore wanted to define the epitopes targeted by vaccine-induced early plasmablast responses. We specifically focused on non-RBD directed neutralizing antibodies and selected for structural characterization PVI.V6-14, an antibody that neutralized the authentic SARS-CoV-2, albeit not very potently, whose amino-acid sequence indicated that it was derived from an unmutated plasmablast.

We determined the structure of PVI.V6-14 in complex with the spike glycoprotein and showed that the antibody targets a hitherto uncharacterized epitope. The antibody bound on the side of NTD, away from RBD, suggesting that the mechanism by which it neutralizes the virus is more complex than simple steric hindrance of ACE2 receptor binding. Our structure showed that PVI.V6-14 stabilized NTD by inserting its HCDR3 loop into a hydrophobic pocket between two beta sheets. Indeed, this stabilization of NTD allowed us to obtain a high resolution cryo-EM reconstruction and trace the entire N-terminal domain polypeptide chain.

Our data show that PVI.V6-14 bound NTD in a configuration that is inconsistent with binding of another NTD antibody class^32^ that allosterically competes with a heme metabolite biliverdin for NTD binding. PVI.V6-14, however, also competes with biliverdin and its neutralization potency is reduced in presence of biliverdin. Whether biliverdin or its metabolites have any biological relevance in the infectious cycle of the virus remains to be explored. A previous study suggested that SARS-CoV-2 coopted biliverdin to evade antibody responses directed to this hydrophobic cavity^32^. Our data, however, show that mRNA SARS-CoV-2 vaccination induced antibody responses to this particular epitope, indicating that, under physiological processes of antigen presentation to B cells, this epitope was, at least partially, unoccupied.

The emergence of viral variants of concern that contain multiple substitutions and deletions in NTD indicates that NTD, like RBD, is under immune pressure. PVI.6-14 binds a relatively conserved epitope since the antibody maintained binding to B.1.1.7 (Alpha) and B.1.351 (Beta) lineages. Binding to P.1 (Gamma), which carries a R190S substitution in this pocket, was in contrast, almost completely lost. However, we have shown earlier that mAb PVI.6-14 also loses neutralizing activity to B.1.1.7 and B.1.351. Should antibodies of this class also be present in the memory B cell compartment, they could potentially return to germinal centers for somatic hypermutation and affinity maturation against P.1 or related viruses with mutations within the hydrophobic pocket to gain breadth and neutralization potency. This is particularly plausible in light of the germline nature of PVI.V6-14. Thus, targeting NTD offers an alternative to RBD-centric vaccine immunogen design^36^ and paves the way to next-generation vaccines that target NTD in addition to RBD and potentially to antibody therapeutics that combine both RBD- and NTD-targeting neutralizing mAbs.

## Methods

### Protein expression and purification

All recombinant proteins were produced using Expi293F cells (Life Technologies). Spike proteins for enzyme-linked immunosorbent assay (ELISA) were cloned into a mammalian expression vector, pCAGGS as described earlier^37,38^ and purified after transient transfections with each respective plasmid. Six-hundred million Expi293F cells were transfected using the ExpiFectamine 293 Transfection Kit and purified DNA. Supernatants were collected on day four post transfection, centrifuged at 4,000 g for 20 minutes and finally, the supernatant was filtered using a 0.22 µm filter. Ni-nitrilotriacetic acid (Ni-NTA) agarose (Qiagen) was used to purify the protein via gravity flow and proteins were eluted as previously described^37,38^. The buffer was exchanged using Amicon centrifugal units (EMD Millipore) and all recombinant proteins were finally re-suspended in phosphate buffered saline (PBS). Proteins were also run on a sodium dodecyl sulphate (SDS) polyacrylamide gels (5–20% gradient; Bio-Rad) to check for purity^39,40^. For cryo-EM, SARS-CoV-2 HexaPro spike was used^41^. The protein was transiently expressed in Expi293F cells (ThermoFisher). Five to 7 days post-transfection, supernatants were harvested by centrifugation and further purified using immobilized metal affinity chromatography (IMAC) with cobalt-TALON® resin (Takara) followed by Superdex 200 Increase 10/300 GL size exclusion column (GE Healthcare).

### Bio-layer interferometry (BLI)

Bio-layer Interferometry (BLI) experiments were performed using the BLItz system (fortéBIO, Pall Corporation). Recombinant SARS-CoV-2 NTD was immobilized on a Ni-NTA biosensor, and mAb PVI.V6-14 was then applied at 2.9 μM to obtain binding affinities and biliverdin was titrated from 60 to 7.7 μM. All measurements were repeated in subsequent independent experiments. K_D_ values were obtained through local fit of the curves by applying a 1:1 binding isotherm model using vendor-supplied software. All experiments were performed in Tris-HCL pH 7.5 and at room temperature.

### ELISA

25ng of the SARS-CoV-2 spike proteins were adhered to high-capacity binding, 96 well-plates (Corning) overnight in phosphate buffered saline (PBS). Plates were blocked with 5% bovine serum albumin (BSA) in PBS containing Tween-20 (PBS-T) for 1hr at room temperature (RT). Blocking solution was discarded and 3-fold dilutions of mAb PVI.V6-14 in PBS were added to wells and incubated for 1hr at RT. Plates were then washed three times with PBS-T. Anti-human IgG-Biotin (Abcam) in PBS-T was added to each and incubated for 1hr at RT. Plates were then washed three times with PBS-T. Streptavidin-horseradish peroxidase (HRP) (Abcam) in PBS-T was added to each and incubated for 1hr at RT. Plates were then washed three times with PBS-T Plates were developed using 1-Step Ultra 3,3’,5,5’-tetramethylbenzidine (TMB) substrate (ThermoFisher), stopped with sulfuric acid and immediately read using a plate reader at 450nm. Data were plotted in Prism 9 (GraphPad Software) and area under the curve (AUC) calculated.

### Neutralization assays

Twenty-thousand cells in 100 µls per well were seeded on sterile 96-well cell culture plates one day prior to the neutralization assay. In general, cells were used at 90% confluency to perform the assay. Serial dilutions of the mAb samples were made in 1X minimal essential medium (MEM; Life Technologies) starting at 30 µg/ml. All work with authentic SARS-CoV-2 (isolate USA-WA1/2020) was done in a biosafety level 3 (BSL3) laboratory following institutional biosafety guidelines and has been described in much greater detail earlier^37,42^. An authentic SARS-CoV-2 isolate (USA-WA1/2020, BEI Resources NR-52281) was pre-incubated with 25 µM of biliverdin for 20 min in 1X MEM. One thousand median cell culture infectious doses (TCID_50_s) of authentic virus were added to each mAb sample (with or without biliverdin) and the mixture was incubated for 1 hour inside the biosafety cabinet. Media from the cells was removed and 120 µls of the virus-mAb (+/-biliverdin) mixture was added onto the cells for 1 hour at 37°C. After one hour, the mixture was removed and 100 µls of each corresponding dilution was added to every well. In addition, 100µls of 1X MEM was also added to every well. Cells were incubated for 48 hours at 37°C after which the media was removed and 150 uls of 10% formaldehyde (Polysciences) was added to inactivate the virus. For assay control, remdesivir was used at 10 µM. After 24 hours, cells were permeabilized and stained using an anti-nucleoprotein antibody 1C7 as discussed in detail earlier^37,43^.

### Cryo-EM sample preparation and data collection

SARS-CoV-2 spike HexaPro was incubated with PVI.V6-14 Fab at 1 mg/mL at a molar ratio of 1.5:1 Fab:Spike for 20 minutes at 4°C. 3 μl aliquots of the spike:Fab complex were applied to UltrAuFoil gold R0.6/1 grids and subsequently blotted for 3 seconds at blot force 3 at 20°C and 100% humidity, then plunge-frozen in 100% liquid ethane using an FEI Vitrobot Mark IV. Grids were imaged on a Titan Krios microscope operated at 300 kV and equipped with a 15 eV energy filter and a Gatan K3 Summit direct detector. 10,690 movies were collected in super-resolution counting mode at 15 e−/pix/s for 4.03 s for a total dose of 50 e-/ Å2/s. Images were collected at a magnification of 81,000, corresponding to a calibrated pixel size of 1.12 Å/pixel, 0.56 Å/pixel super-resolution with a defocus range of –2.5 to –0.8 μm.

### Cryo-EM data processing

Data processing was done using Relion^44^. 9,367 micrographs were aligned, and dose weighted using Relion’s implementation of MotionCorr2^45^. The contrast transfer function estimation was calculated using GCTF^46^. Particles were picked with Topaz^47^ with a model trained with a subset of refined, classified particles picked using crYOLO^48^ with particle diameter value of 330Å. 759,324 picked particles were binned to ∼12 Å/pixel and subjected to a 2D classification. 179,294 selected particles were then extracted to ∼6 Å/pixel, subjected to a second round of 2D classification. 88,192 particles were selected were then subjected to one round of 3D classification using a previously processed spike:Fab complex as reference, yielding 60,639 particles in the final subset. Particles were then extracted to 1.12 Å/pixel and aligned using 3D auto-refine then un-binned to 0.56 Å/pixel for final refinement. Un-binned particles were aligned using 3D classification with 1 class and a regularization parameter of 25. Particles were then focus aligned using 3D classification with 1 class, first with a mask encompassing the S1 region for the monomer with the best Fab density, then continued with a mask encompassing the NTD and Fab variable regions. Full spike and focus classified maps were then sharpened using DeepEMhancer.

### Model building and refinement

For the NTD, a published crystal structure (PDB ID 7L2C) was docked into the sharpened focused map in UCSF Chimera^49^ then manually fit using COOT^50^. Previously unbuilt regions were manually built. For the Fab, the variable regions of the structures with the highest sequence identity to PVI.V6-14 (PDB ID 6XWD and 5XI5 for heavy and light chains, respectively) were docked into the focused map in UCSF Chimera then used as a reference for manual building. For the remainder of the spike, a previously published structure (PDB ID 7NTC) was docked into the sharpened full map in UCSF Chimera then manually fit in COOT. All models were then refined in Phenix^51^ using Real-space refinement against their relative maps.

## Acknowledgments

We would like to thank Aaron G. Schmidt for critical reading of the manuscript and Kevin R. McCarthy for help with curating the sarbecovirus spike sequences. We would like to thank the study participants for their generosity and willingness to participate in longitudinal COVID19 research studies. None of this work would be possible without their contributions. We would like to thank Dr. Randy A. Albrecht for oversight of the conventional BSL3 biocontainment facility, which makes our work with live SARS-CoV-2 possible. Part of this work was done at LBMS in Brookhaven. The Laboratory for BioMolecular Structure (LBMS) is supported by the DOE Office of Biological and Environmental Research (KP160711). This work was partially funded by the NIAID Collaborative Influenza Vaccine Innovation Centers (CIVIC) contract 75N93019C00051. This work was partially funded by the Centers of Excellence for Influenza Research and Surveillance (CEIRS, contract # HHSN272201400008C), and by the generous support of the JPB Foundation and the Open Philanthropy Project (research grant 2020-215611 (5384), and by anonymous donors

## Author contributions

A.H.E., V.S., F.K., and G.B. designed the study; C.G.A., D.C.A., J.M.C., I.A.S. and F.A. performed research; C.G.A., D.C.A., J.M.C., A.H.E., V.S., F.K., and G.B. analyzed data; C.G.A. and G.B. prepared the figures; G.B. wrote the paper. All authors edited and commented on the paper.

## Materials & Correspondence

All data are provided in the Supplementary Materials. Requests for material should be addressed to Goran Bajic. The EM maps have been deposited in the Electron Microscopy Data Bank (EMDB) under accession codes: EMD-24402 and EMD-24403 and the accompanying atomic coordinates in the Protein Data Bank (PDB) under accession codes: 7RBU and 7RBV.

## Figures

**Figure S1.**
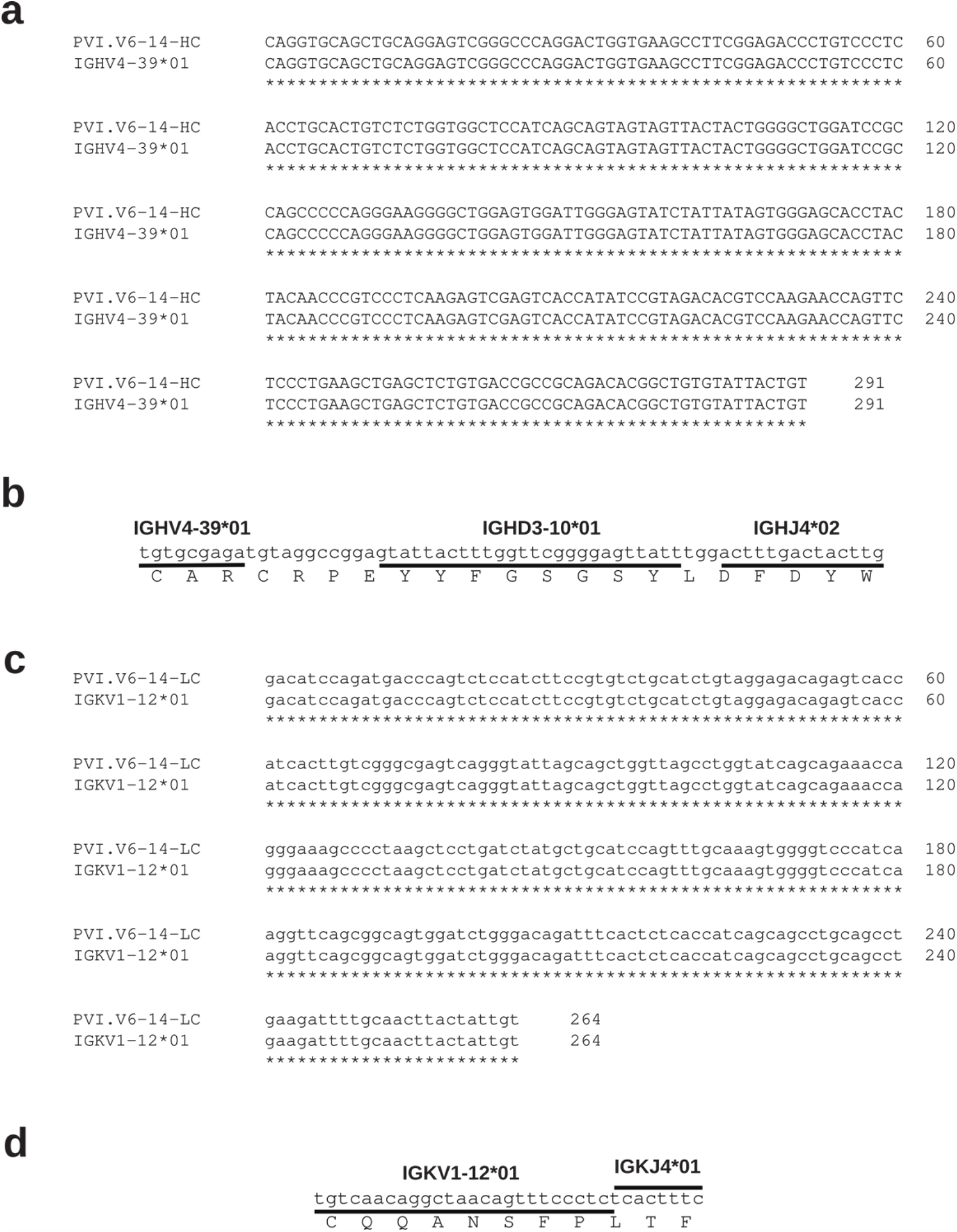
PVI.V6-14 is an unmutated rearranged V(D)J sequence. **(a)** Nucleotide sequence alignment of PVI.V6-14 variable heavy gene (IGHV) with its germline IGHV4-39*01. **(b)** *V, D* and *J* gene fragments mapped onto the nucleotide sequence of PVI.V6-14 HCDR3 loop. The translated amino-acid sequence is shown below. **(c)** Nucleotide sequence alignment of PVI.V6-14 variable kappa light gene (IGKV) with its germline IGKV1-12*01. **(d)** *V* and *J* gene fragments mapped onto the nucleotide sequence of PVI.V6-14 LCDR3 loop. The corresponding amino-acid sequence is shown below.

**Figure S2.**
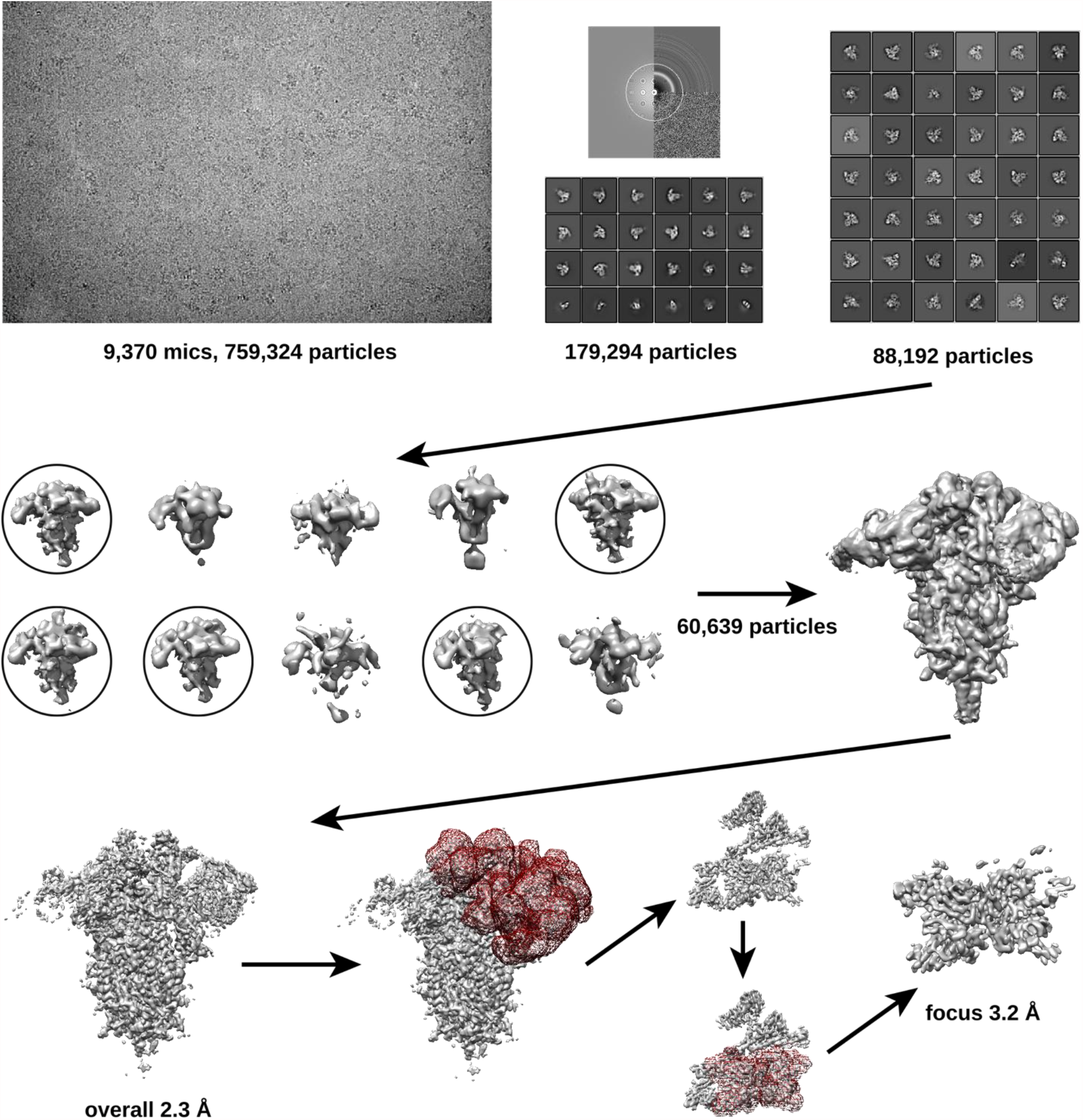
Cryo-EM data processing scheme for PVI.V6-14 Fab bound with SARS-CoV-2 spike. Representative micrograph and its CTF are shown together with two rounds of 2D classification in the top row. The 3D classification and refinement logic tree are shown below. See the Methods section for more details.

**Figure S3.**
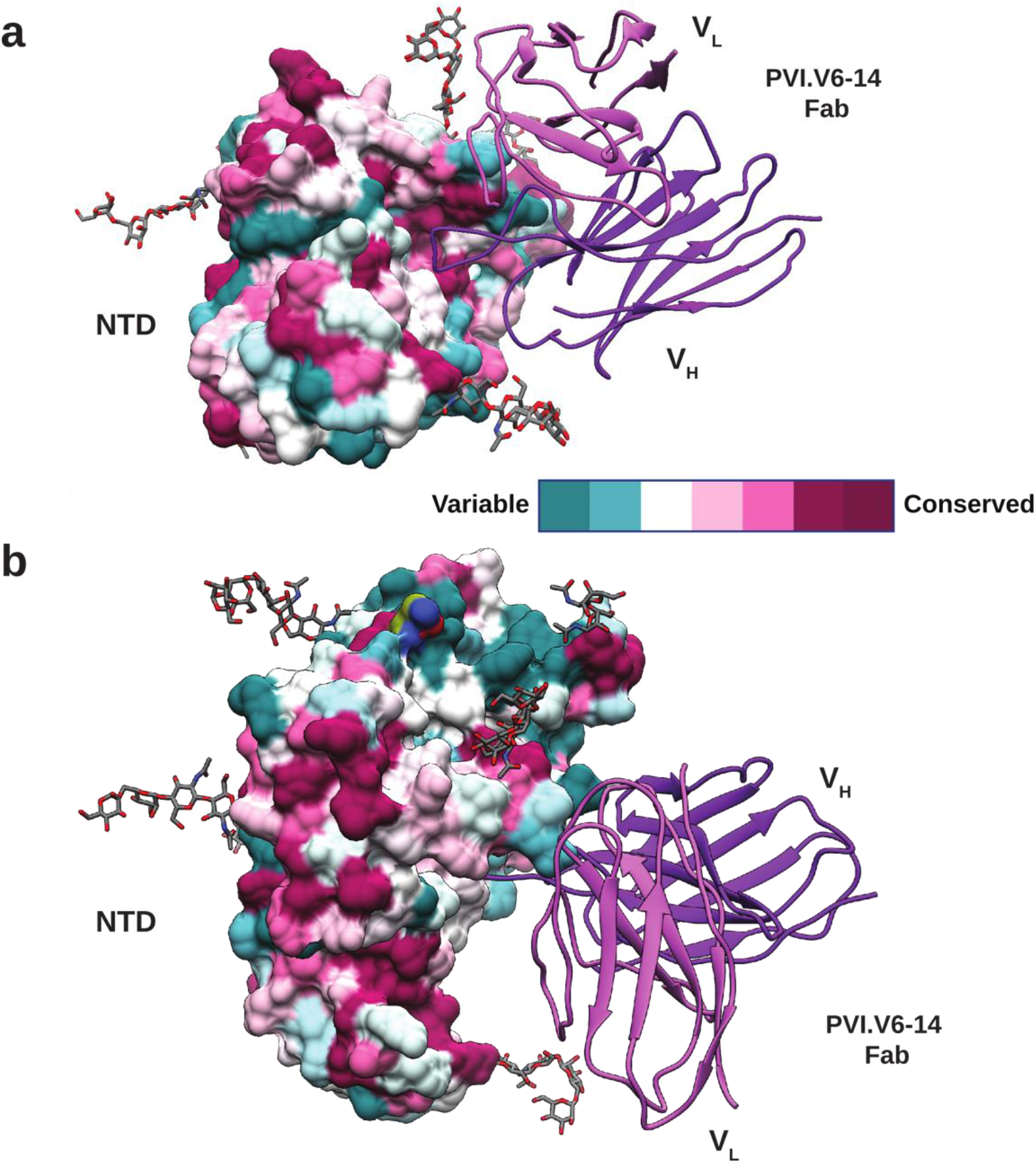
NTD conservation map. Amino-acid sequence conservation of a representative set of spike sequences from the sarbecovirus subgenus aligned and mapped onto the structure of the NTD:PVI.V6-14 complex. The hydrophobic cavity contains conserved residues. Accession codes used: NCBI: KY417152, KY417151, MK211376, KC881006, KF367457, KT444582, KY417150, NC_004718, AY304486, AY304488, AY572034; GISAID Epi-Cov: EPI_ISL_412860, EPI_ISL_1699444, EPI_ISL_1699443, EPI_ISL_1699446, EPI_ISL_1699445, EPI_ISL_804222, EPI_ISL_410540, EPI_ISL_412977, EPI_ISL_402125, EPI_ISL_402131, EPI_ISL_1699447, EPI_ISL_410542, EPI_ISL_410541, EPI_ISL_410544, EPI_ISL_410538, EPI_ISL_1699448, EPI_ISL_410543, EPI_ISL_1699449, EPI_ISL_410539, EPI_ISL_410721

